# Cellular tolerance at the μ-opioid receptor is phosphorylation dependent

**DOI:** 10.1101/252387

**Authors:** Seksiri Arttamangkul, Daniel A Heinze, James R Bunzow, Xianqiang Song, John T Williams

## Abstract

The role of phosphorylation of the μ-opioid receptor (MOR) in desensitization, internalization and long-term tolerance was examined in locus coeruleus (LC) neurons. Viral expression of wild type (exWT) and mutant MORs, where all phosphorylation sites on the C-terminus (Total Phosphorylation Deficient (TPD)) were mutated to alanine, were examined in a MOR knockout rat. Both expressed receptors acutely activate potassium conductance similar to endogenous receptors in wild type animals. The exWT receptors, like endogenous receptors, displayed signs of tolerance after chronic morphine treatment. There was however a loss of agonist-induced desensitization and internalization in experiments with the TPD receptors. In addition the development of tolerance was not observed in the TPD receptors following chronic morphine treatment. The results indicate a key role of C-terminal phosphorylation in the expression of acute desensitization, trafficking and long-term tolerance to morphine.

## Introduction

Considerable effort has been aimed at characterizing the mechanisms that underlie acute μ-opioid receptor (MOR) dependent desensitization and cellular tolerance (Williams et al., 2013). One key step thought to be important in these processes is the phosphorylation of sites on the cytoplasmic loops and C-terminal tail following receptor activation. On the C-terminal tail of the MOR there are 11 possible phosphorylation sites. Two specific cassettes, amino acid residues 354 to 357 (TSST) and 375 to 379 (STANT), are phosphorylated following application of agonists that induce desensitization and internalization (Lau et al., 2011). Point mutations of serine (S) and threonine (T) residues in the STANT sequence resulted in a decrease in agonist induced arrestin recruitment and internalization, but do not eliminate the induction of acute desensitization (Lau et al., 2011; Birdsong et al., 2015). Complete alanine mutation of both the TSST and STANT sequences significantly reduced, but did not completely eliminate acute desensitization. Although the TSST and STANT sequences are known to be agonist-dependent phosphorylation sites, there are four other serine and threonine sites on the C-terminus that are either phosphorylated constitutively or by agonist dependent kinases (Williams et al., 2013). It is not known if phosphorylation of these sites alters the regulation of MORs.

The present study investigated the role of C-terminus MOR phosphorylation on acute signaling, desensitization, internalization and cellular tolerance in rats. The desensitization of MOR in rat locus coeruleus neurons has been studied extensively in wild type animals. In order to study wild type and mutant MORs, a MOR-knockout rat model was used and MORs were virally expressed. Studying receptor trafficking of expressed receptors was enabled by linking a green fluorescence protein (GFP) to the N-terminus of the MOR constructs. Two versions of MOR were expressed, wild-type (exWT) and total phosphorylation deficient receptors (TPD) where all 11 possible phosphorylation sites on the C-terminus were mutated to alanine. The results show that both receptors activate a hyperpolarizing potassium current. There was no significant difference in the kinetics of activation between WT, exWT, and TPDs. Thus acute activation of virally expressed MORs was not significantly different from endogenous receptors. There was however a near complete loss of desensitization, internalization, and long-term tolerance in neurons expressing TPD receptors. This study demonstrates a key role of phosphorylation in the both acute- and long-term actions of opioids on single neurons.

## Results

### MOR Knockout

Recordings from LC neurons in brain slices from the MOR knockout animal confirmed that there was no current or hyperpolarization induced by opioids with no detectable difference in the activation of alpha-2-adrenoceptors, orphanin FQ, or M3-muscarine receptors (Figure 1, S1).

**Figure 1.**
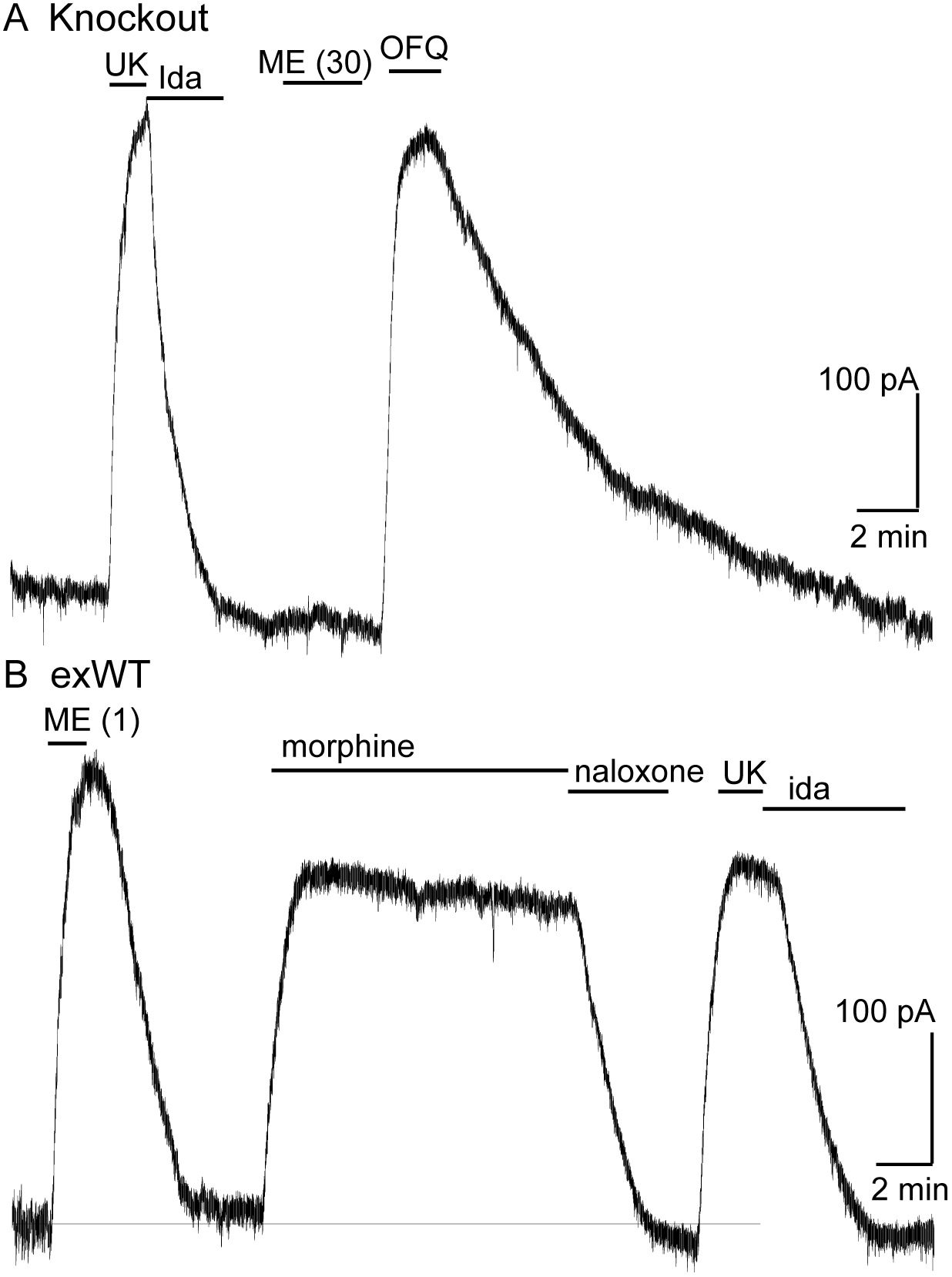
Locus coeruleus neurons are not sensitive to opioids in the MOR knockout rat. A, from a MOR knockout animal where the alpha-2-adrenoceptor agonist, UK14304 (3 μM) and OFQ both activate potassium currents whereas ME (30 μM) had no effect. B, from a neuron where the wild type MOR (exWT) was expressed in the MOR knockout animal. In this recording ME (1 μM), morphine (10 μM) as well as UK14304 (3 μM) all caused outward currents.

### MOR expression

Microinjections of adeno-associated virus type 2 containing either the exWT or TPD receptors were made bilaterally into the LC and after 2-4 weeks the GFP tagged MORs were visualized with an Olympus macroview microscope (Figure 2A). Images obtained with a 2-photon microscope showed green fluorescence on the plasma membrane and in the cytoplasm of LC neurons. Plasma membrane receptors were selectively targeted and labeled using an anti-GFP nanobody conjugated to Alexa594 dye (Figure 2B-D). There was no labeling of cells from animals that did not express GFP tagged MORs (Figure 2E,F).

**Figure 2.**
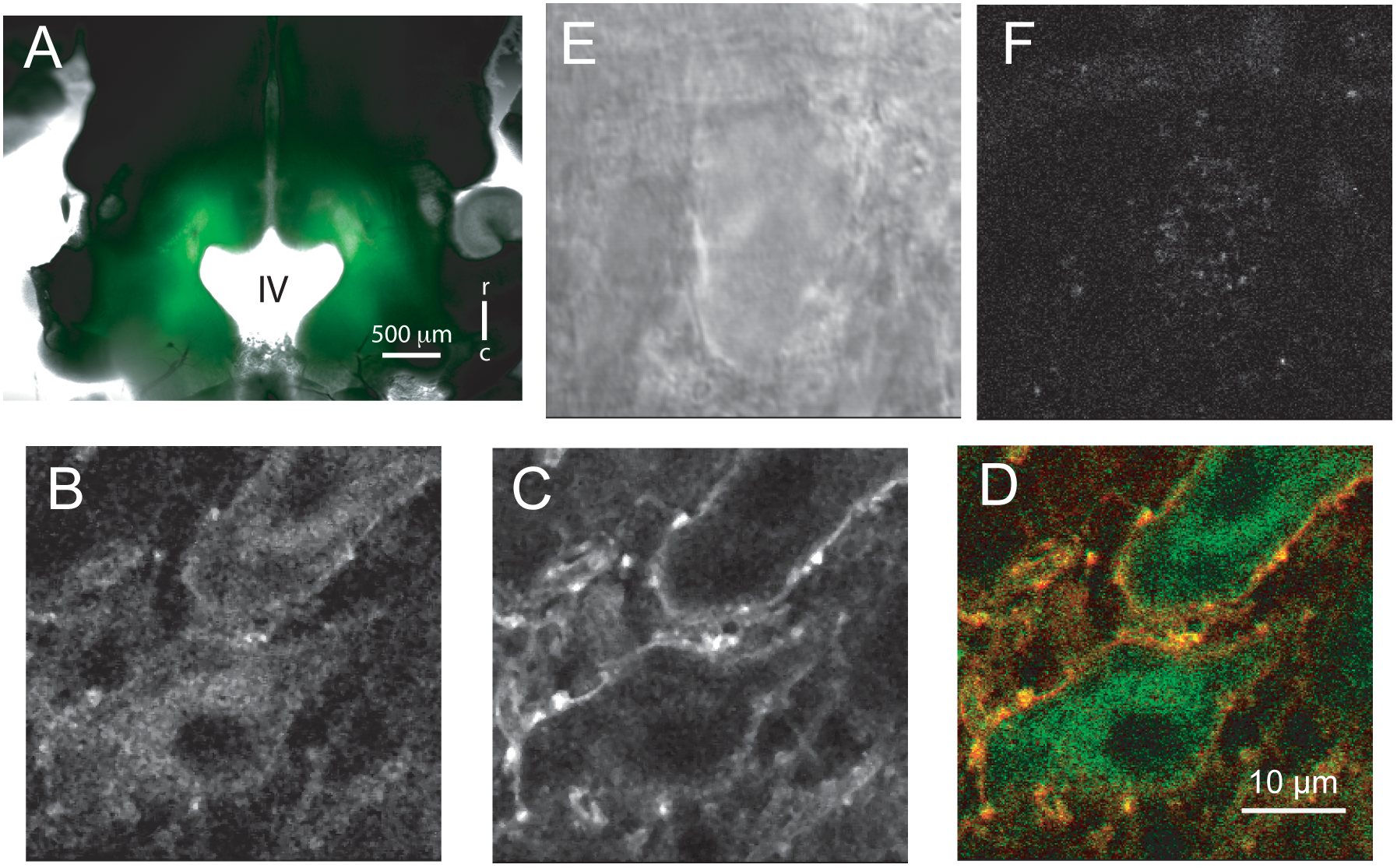
Microinjection of virus expressing GFP-tagged exWT receptors in the locus coeruleus of MOR knockout rat. A, low power image of the GFP fluorescence observed in a horizontal slice containing the LC. B, shows an image obtained with a 2-photon microscope showing the GFP fluorescence. C, the same neuron showing the plasma membrane associated receptors following incubation of the slices with an anti-GFP nanobody conjugated with alexa594. D, the merged image of the GFP and alexa594 fluorescence. E, a scanning DIC image of a LC neuron from a knockout animal. F, the same neuron following incubation of the slice with the anti-GFP nanobody conjugated with alexa594. Scale bar = 10 μm.

### Electrophysiology of the virally expressed receptors

Recordings were made from slices from animals 2-4 weeks following viral microinjection. Whole-cell voltage clamp or intracellular membrane potential recordings were made from LC neurons. In each case, application of [Met^5^]enkephalin (ME, 300 nM, an EC50 concentration in slices from wild type animals) was used as an estimate of the level of receptor expression. Cells where the outward current or hyperpolarization induced by this application was either very large or small were not included in the study.

The kinetics of receptor-dependent activation of G protein-gated inwardly rectifying potassium (GIRK) conductance was examined with the photolysis of caged-enkephalin (CYLE) using whole-cell recordings in the voltage clamp configuration (Figure 3). The rate of activation (10-90%, exWT 238±33 ms, n=9, TPD 265±39 ms, n=19) and the time to the peak of the outward current (exWT 1.78±0.12 s, n=12, TPD 2.05±0.20, n=18) were the same between exWT and TPD and similar to those from experiments made from cells expressing wild type receptors (Figure 3). Likewise the return to baseline following photolysis of the caged antagonist naloxone (CNV-Nal) in the presence of ME (1 μM) was not different in recordings made from the two receptors (exWT 1.57±0.08 s, n=7, TPD 1.53±0.15, n=6, p>0.05). When the high affinity agonist endomorphin-1 (100 nM) was used, the time constant of CNV-Nal-induced inactivation in exWT receptors (4.06±0.41 s, n=8) was not significantly different from that measured with TPD receptors (5.37±0.59 s, n=10, p>0.05). The current amplitudes induced by endomorphin-1 (290±37 pA in exWT and 255±44 pA in TPD) were also similar (p>0.05). Thus, in spite of the multiple mutations along the C-terminus and the presence of GFP at the N-terminus, the acute signaling of virally expressed TPD and exWT receptors was not significantly different compared to receptors from wild type animals.

**Figure 3.**
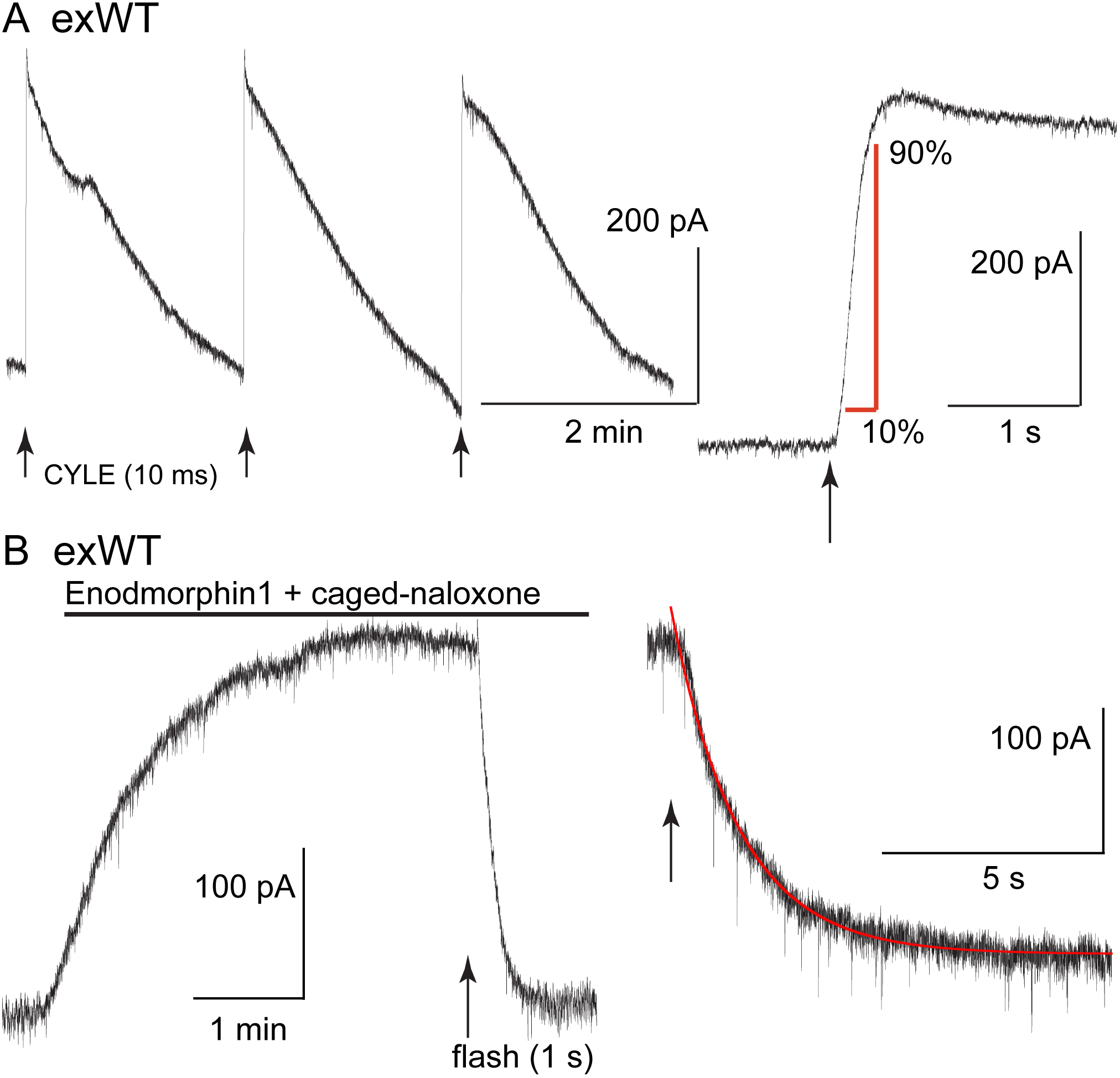
The kinetics of activation and inhibition of expressed wild type receptors (exWT) in LC neurons from the MOR knockout measured with photolysis of caged enkephalin (CYLE). A, repeated photolysis events (1/2 min) resulted in rapidly rising outward currents. Right side, the rate of rise was measured as the time it took to go from 10 to 90% of the peak. B, the high affinity agonist endomorphin 1 (100 nM) was applied along with caged-naloxone (5 μM). At the arrow a 1 s photolysis flash was applied and the decrease in the outward current was measured. Right side is an expanded time base of the decrease in outward current and the single exponential fit to that decline.

### Desensitization

Acute receptor desensitization was compared in slices from animals injected with either exWT or TPD receptors. Two measures of desensitization were made, the decrease in the peak (hyperpolarization for intracellular recordings or current for whole-cell recordings) during the application of a saturating concentration of ME (30 μM, 10 min) and the relative decrease in response to an EC50 concentration of ME (300 nM) following the washout of the saturating concentration of ME (Figure 4, S4). The results from experiments with exWT receptors were similar to results from wild type animals that have been published previously. During the application of ME (30 μM) the membrane potential hyperpolarization measured with intracellular recording declined to 75±5% of the peak (n=6). The current measured under voltage clamp with whole-cell recording declined to 54±4% of the peak (n=12). Following the washout of saturating ME (30 μM), the hyperpolarization induced by EC50 ME (300 nM) was reduced to 30±2% (n=5) and the current to 23±5% (n=10) of the pre-desensitization response. Following a 20-30 min wash of ME (30 μM) the application of ME (300 nM) approached the pre-desensitization control with whole cell voltage clamp or intracellular recordings.

**Figure 4.**
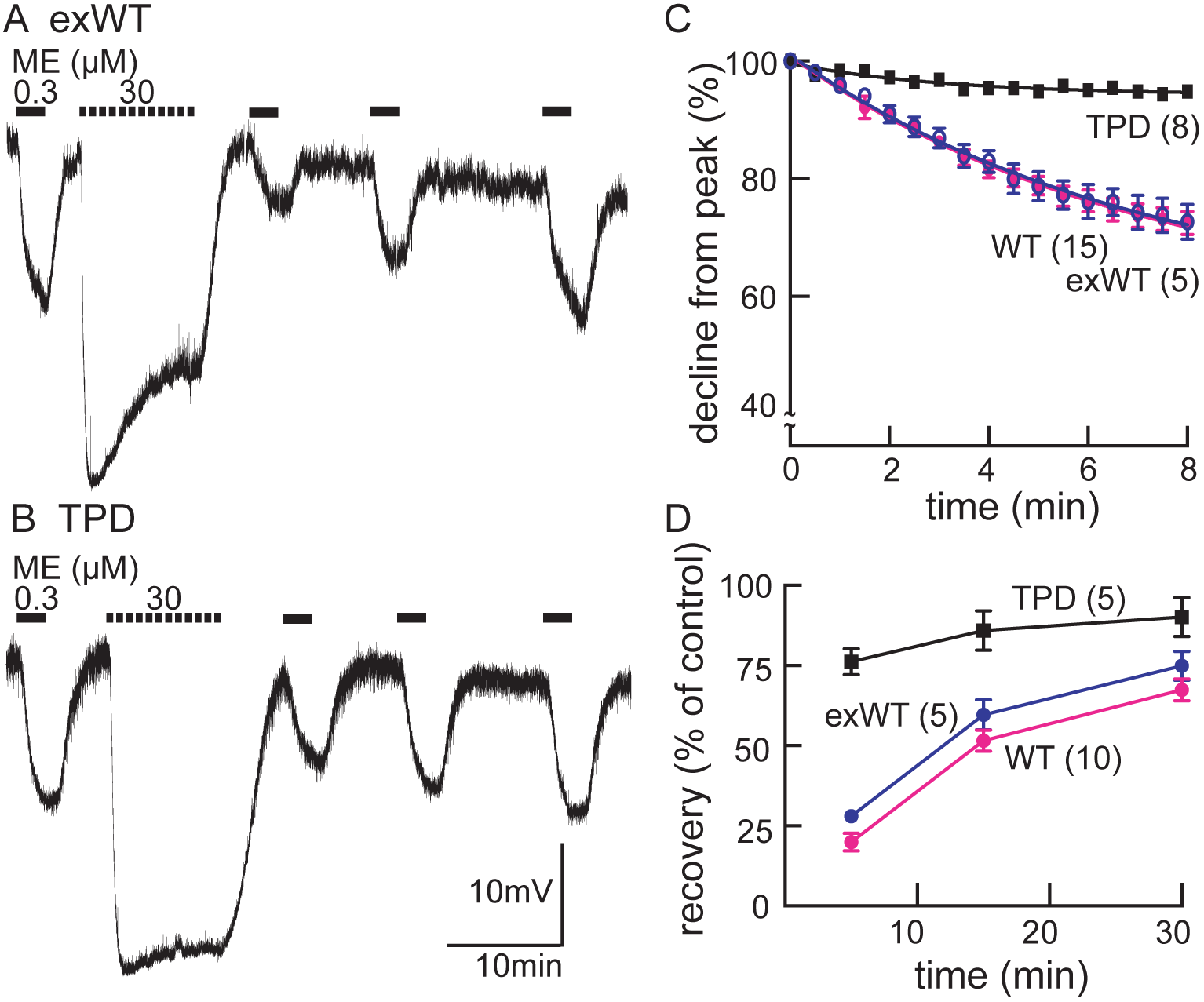
Desensitization is largely blocked in experiments carried out in neurons expressing the TPD. Experiments carried out with intracellular recordings of membrane potential. A, an example of an experiment with a cell expressing the exWT receptor. An EC50 was applied before and following application of a saturating concentration of ME. The amplitude of the hyperpolarization induced by ME (30 μM) decreased during the 10 min application. The hyperpolarization induced by ME (300 nM) was reduced and recovered slowly following washout of ME (30 μM). B, the same experiment carried out in neurons that expressed the TPD receptor. C, summarized results showing the decline in the hyperpolarization during the application of ME (30 μM). There is only a small decline in experiments carried out with the TPD receptor. D, summarized results showing the recovery from desensitization. Experiments with neuron in wild type animal (WT) and the expressed exWT receptors show a slow recovery, whereas there was little sign of desensitization in the experiments from the TPD receptor.

The results obtained with the TPD receptors were significantly different. With intracellular recordings, the hyperpolarization was reduced to 94±1% (n=7, p=0.0003) of the peak and the current declined to 80±2% of the peak (n=13, p=0.003) in whole cell recordings. Likewise there was only a small decrease in the response to ME (300 nM) following washout of the saturating concentration of ME (76±4%, n=5, p=0.0001 in intracellular recording and 81±9% of pre-desensitization, n=11, p=0.0005 in voltage clamp). Thus by two measures obtained under two recording conditions, acute desensitization was markedly reduced in neurons expressing TPD receptors.

### Protein kinase C dependent desensitization

MOR desensitization induced by protein kinase C (PKC) has been proposed to be the result of phosphorylation at S363, T370 or S/T356-357 (Wang et al., 2002; Feng et al., 2011; Chen et al., 2013; Illing et al., 2014). In previous work using wild type animals, receptor desensitization was augmented by manipulations that increased the activity of PKC. The PKC activators, phorbol 12,13-dibutyrate and phorbol 12-myristate 13-acetate increased the decline from the peak hyperpolarization during the application of ME (30 μM, 10 min) and caused a small reduction of the hyperpolarization induced by ME (300 nM). Muscarine, thought to activate PKC by a Gq-dependent mechanism, also induced a large increase in apparent desensitization (Arttamangkul et al., 2015). In intracellular recordings from both exWT and TPD receptors, muscarine facilitated the desensitization induced by ME (30 μM, Figure 5). In the presence of muscarine, the initial amplitude of the hyperpolarization induced by ME (300 nM) decreased by 43±8% in experiments from exWT, n=5 and by 49±8% from TPD, n=6. The decline from the peak hyperpolarization induced by ME (30 μM, 10 min) was also facilitated (Figure 5). Although the decline from the peak in experiments examining the TPD receptors was small, it was over double that measured in the absence of muscarine (6±1%, n=7 *vs.* 15±2% in muscarine, n=6, p=0.0009, unpaired two tailed T-test). Likewise, in whole-cell voltage clamp experiments, the current amplitude evoked by photolysis of CYLE was significantly decreased by muscarine and returned to baseline following the application of the muscarinic antagonist scopolamine (Figure S5). Thus the presumed activation of PKC induced by muscarine acts by a mechanism that is independent of phosphorylation of the C-terminal of MOR.

**Figure 5.**
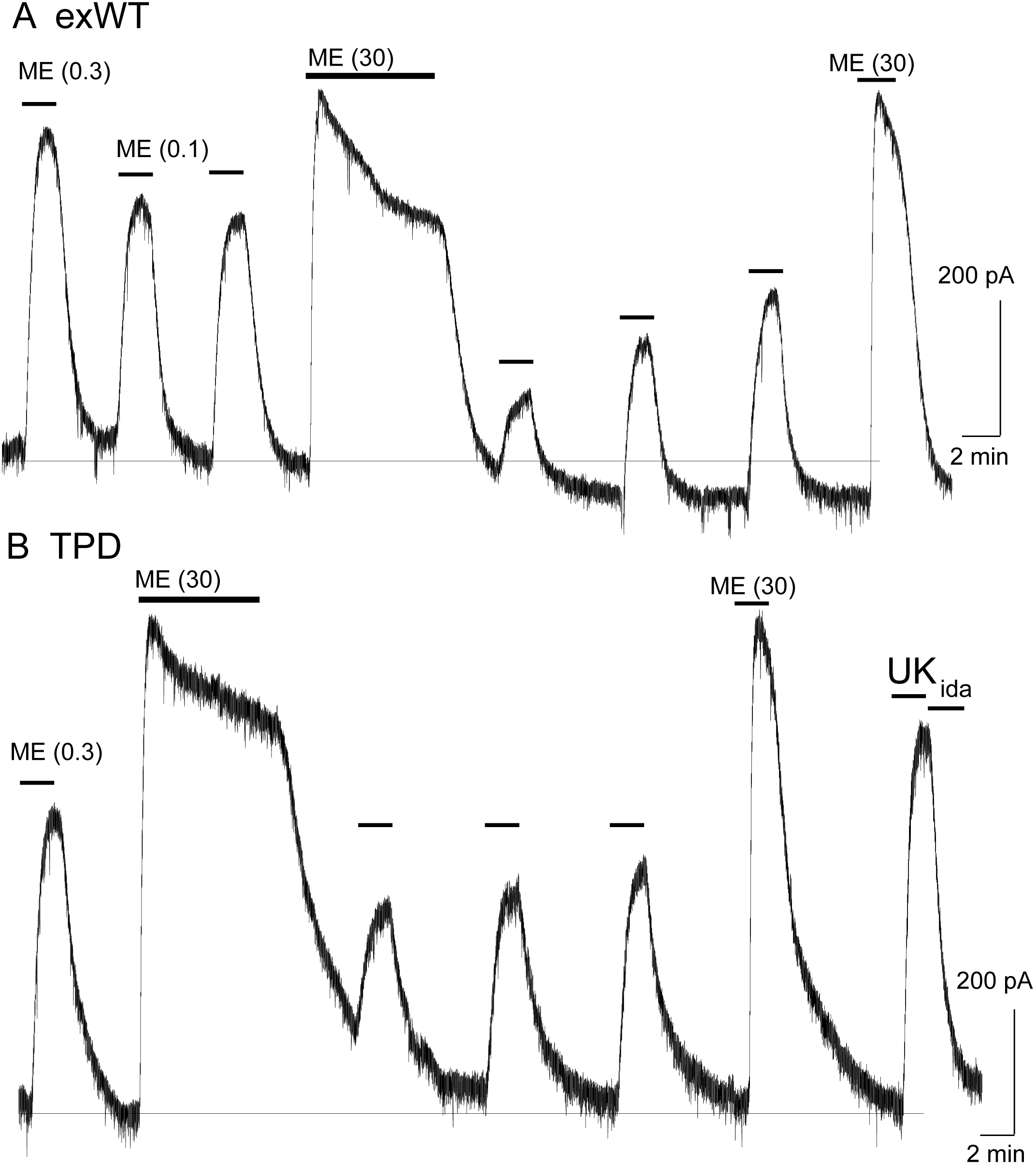
Muscarine inhibits ME-induced hyperpolarization and increases desensitization of WT, exWT and TPD receptors. Recording of membrane potential made using intracellular electrodes. A, a recording from a neuron in a slice from a wild type animal. The hyperpolarizations induced by ME (300 nM) and noradrenline (NA 5 μM, plus cocaine 3 uM and prazosin 500 nM to prevent reuptake) were both decreased in the presence of muscarine (10 μM). The hyperpolarization induced by ME (30 μM) peaked and declined during the 10 min application. B, the same experiment carried out in a recording from an exWT receptor. The hyperpolarization induced by ME (300 nM) was reduced and the decline in the hyperpolarization induced by ME (30 μM) was increased in the presence of muscarine (10 μM). C, an experiment with the TPD receptor showing the lack of decline in the hyperpolarization during the application of ME (30 μM). D, the same experiment carried out in the presence of muscarine (10 μM). The initial hyperpolarization induced by ME (300 nM) is reduced and there is a greater decline in the hyperpolarization during the application of ME (30 μM). E, summary of the inhibition of the initial hyperpolarization induced by ME and NA in WT, exWT and TPD receptors, two-tailed paired t-test (** p<0.01, ****p<0.0001). F, summary of the decline in the hyperpolarization in control and in the presence of muscarine. The muscarine-induced increase in decline in experiments from WT and exWT receptors is the same (WT n=15 in control 11 in muscarine; exWT n=5 in control, 5 in muscarine). There is also a small but significant increase in the decline found in experiments with the TPD receptor (n=8 in control, 6 in muscarine, p<0.0001, two way ANOVA, Bonferroni)

### Receptor trafficking – acutely and following chronic morphine treatment

Receptor internalization was studied by immuno-labeling the extracellular N-terminal GFP of plasma membrane-associated exWT and TPD receptors. Living slices were incubated in an anti-GFP nanobody-Alexa594 for 30-45 min, placed in a superfusion chamber and visualized with 2-photon microscopy (Figure 6). Labeled receptors were imaged before the application of a saturating concentration of ME (30 μM, 10 min) and at the end of the 10 min application. Similar to a previous study in mouse LC neurons (Arttamangkul et al., 2008), this treatment induced the internalization of exWT receptors (Figure 6A top). When TPD receptors were examined using the same treatment protocol there was no detectable internalization (Figure 6A bottom). Thus, ME induced internalization of TPD receptors was completely disrupted (exWT receptor internalization was 38±3% of the total fluorescence, n=7; internalization of the TPD was 10±5% of the total fluorescence, n=6, p=0.0004).

**Figure 6.**
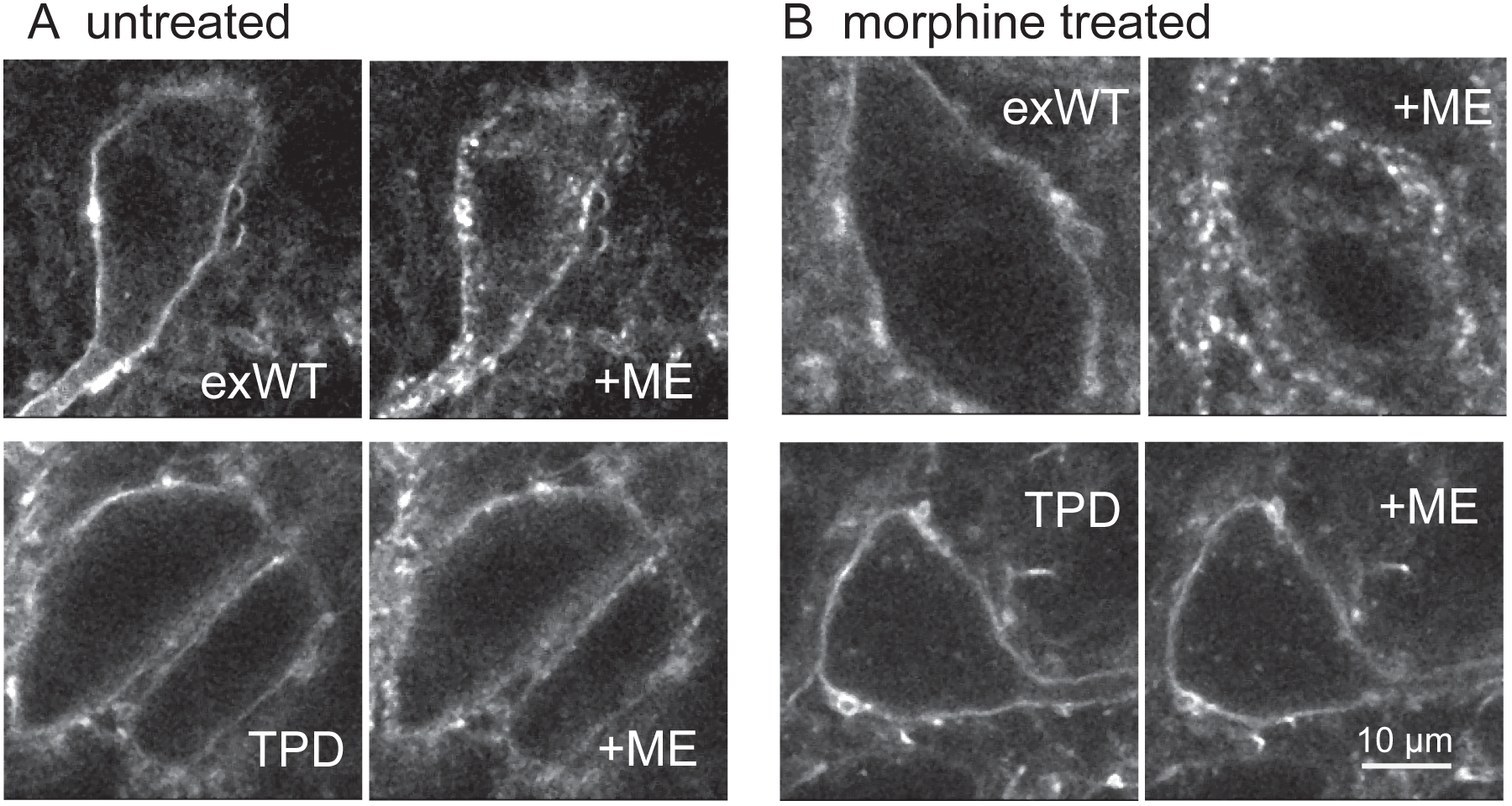
Receptor trafficking of the TPD is blocked in slices from untreated and morphine treated animals. Images from exWT (top) and TPD (bottom) showed receptor distribution before and following application of ME (30 μM, 10 min). The exWT receptors became punctate and moved into the cytoplasm in slices from untreated (A, top) and morphine treated (B, top) animals. The TPD receptors did not traffic in slices from untreated (A, bottom) or morphine treated animals (B, bottom). Scale bar = 10 μm.

The development of long-term tolerance induced by chronic morphine treatment after expression of exWT and TPD receptors was examined next. A previous study in mouse LC found that the recycling of Flag-MORs was increased after chronic morphine treatment in the arrestin3 knockout animals (Quillinan et al., 2011). One possibility is that chronic morphine treatment may result in the modulation the trafficking pathway of expressed exWT and TPD receptors. Animals were microinjected with virus to express either exWT or TPD and after 7-10 days treated chronically with morphine (80 mg/kg/day) using osmotic mini pumps. After 6 or 7 days brain slices were prepared in morphine-free solutions and slices were prepared for 2-photon imaging. As expected, exWT receptors were internalized in slices from morphine treated animals (49±6%, n=4, Figure 6 B top). There was however, no internalization of TPD receptors in slices from morphine treated animals (n=4, Figure 6B bottom). Thus the inability to induce internalization of TPD receptors was maintained following chronic morphine treatment.

### Tolerance following chronic morphine treatment

In whole-cell voltage clamp experiments using slices from morphine treated animals that expressed exWT receptors, a saturating concentration of ME (30 μM) resulted in an outward current that peaked and declined to 41±2% (n=8) of the peak. The current induced by a subsequent application of ME (300 nM) was significantly depressed relative to the control (Figure 7B). Unlike the results obtained in slices from untreated animals, the current induced by ME (300 nM) never recovered completely over a period of 20-30 min after washout of the saturating concentration. Thus as has been reported previously in experiments using wild type animals that were chronically treated with morphine, desensitization was augmented and the recovery from desensitization was reduced.

**Figure 7.**
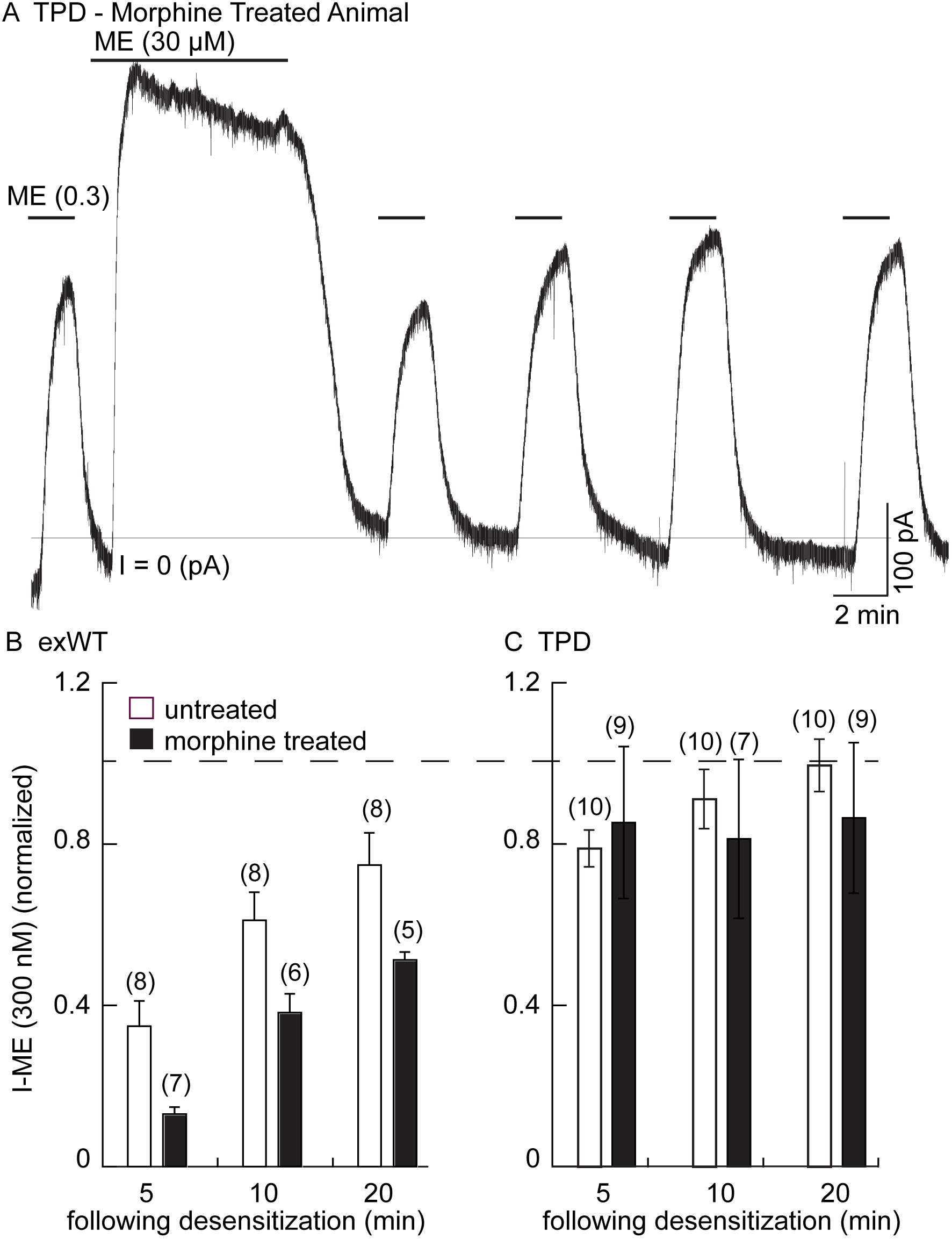
There is no sign of long-term tolerance induced in the TPD receptor following chronic morphine treatment. Whole cell voltage clamp experiments. A, an example from a morphine treated animal expressing the TPD receptor. There was a small decline in the current induced by a saturating concentration of ME (30 μM, 10 min) and a small and transient decrease in the current induced by ME (0.3 μM) following the washout of the ME (30 μM). B, summary shows the recovery from desensitization induced by ME (30 μM, 10 min) in exWT. There is significantly less recovery from desensitization seen in experiments from morphine treated animals than untreated controls with the exWT receptor (p=0.0001, two way ANOVA, Bonferroni). C, summarized results from untreated and the morphine treated animals expressing the TPD receptors. There was a small (not significant) decrease in the ME (0.3 μM) current immediately after the ME (30 μM) treatment in experiments from animals expressing the TPD receptor that did not change with time. There were no difference between the recoveries from the untreated and morphine treated animals at any points (P>0.3, ANOVA Bonferroni).

In experiments from morphine treated animals expressing the TPD receptor, there was no significant change in any measure of opioid action. The decline from the peak current induced by a saturating concentration of ME (30 μM 10 min) was 83±4% in TPD (n=8) and not different from experiments in slices taken from untreated animals (80±2%, n=13, p>0.05, unpaired T-test). The current induced by ME (300 nM) following washout of the ME (30 μM) solution was also not different from slices of untreated animals (Figure 7C).

Finally, the initial rise in current induced by photolysis of CYLE was not different in slices from untreated and morphine treated animals (10-90% rise time of the first flash, untreated exWT 255±35 ms (n=10), TPD 219±32 ms (n=14); morphine treated exWT 201±21 ms (n=10), TPD 211±23 (n=10). Thus the rising phase of the current was not changed following chronic morphine treatment. Another measure of desensitization was examined in slices from morphine treated animals using photolysis of caged-enkephalin (Figure 8). The peak current and 10-90% rise time was measured in response to repeated 10 ms flashes before and after a longer 100 ms flash. In experiments from exWT expressing cells the amplitude of the current declined significantly by the 5^th^ flash. Following the long flash, there was a step decrease in the peak current measured in the exWT receptors (Figure 8C). Likewise the rise time was significantly slowed (Figure 8D). In contrast, the long flash did not alter the peak amplitude or activation rate of the 10 ms flash-induced currents in recordings from cells expressing TPD receptors. Thus, the normalized peak currents and activation rates following the long flash were significantly different between exWT and TPD receptors further indicating that TPD receptors display little to no desensitization even in slices from morphine treated animals.

**Figure 8.**
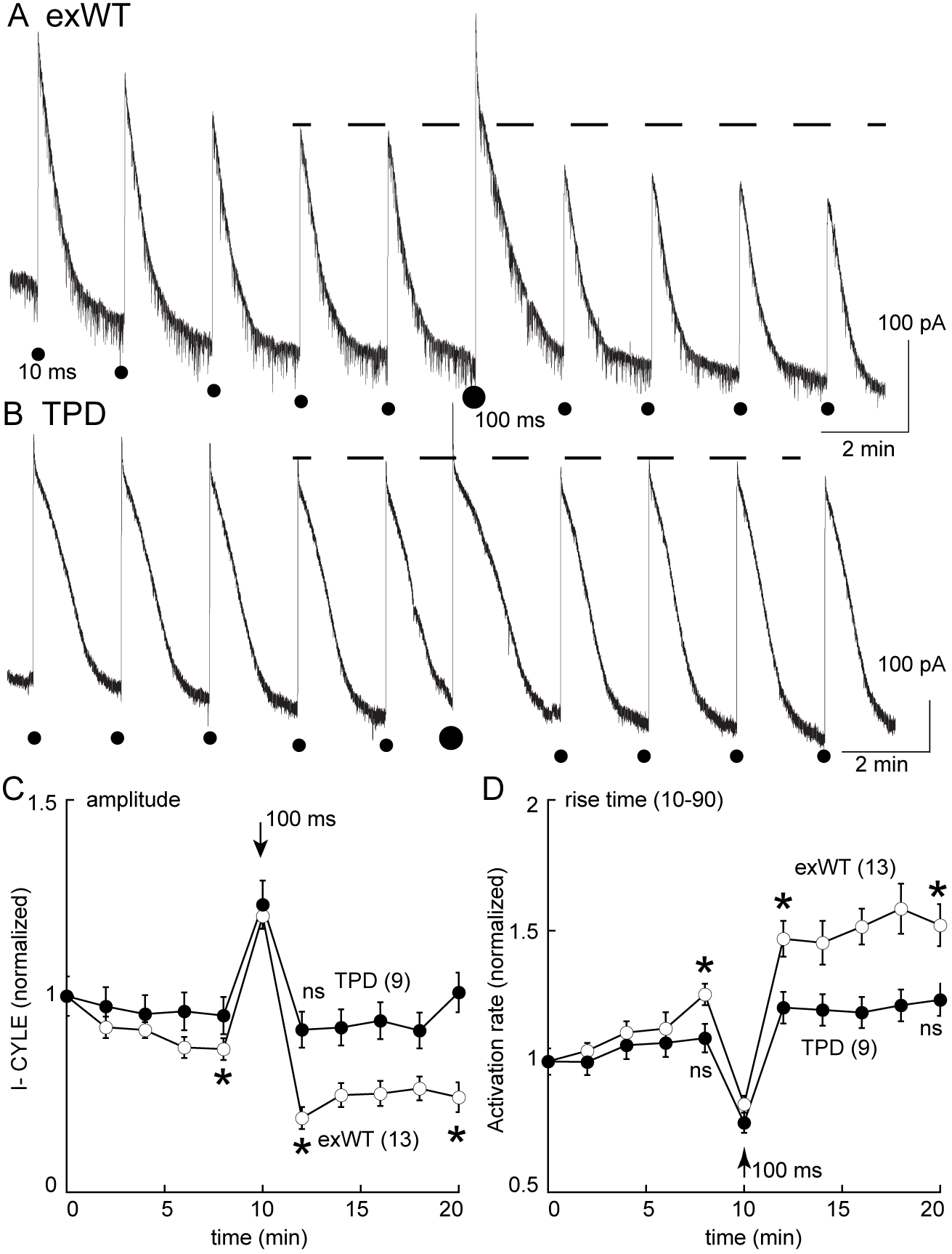
Measures of acute desensitization induced by photolysis of caged enkephalin (CYLE) in slices from morphine treated animals. Whole cell voltage clamp recordings were made from neurons expressing the TPD or exWT receptors. A, exWT and B, TPD are example experiments where photolysis of CYLE was carried out every 2 min. The initial duration of the flash was 10 ms, after 5 flashes the duration of the flash was increased to 100 ms and subsequent flashes were 10 ms. C, shows summarized results of the amplitude of the outward currents. In recordings from exWT receptors the amplitude of the current induce by the 10 ms flash decreased steadily (p=0.002). Increasing the duration of the flash increased the peak current and the amplitude of the subsequent currents induced by 10 ms flashes was decreased (p<0.0001). The current induced by the 10 ms flashes in experiments from the TPD receptor were not changed (p>0.3). The decline in current in the exWT receptors is significantly different than in the TPD receptors (p=0.0001, ANOVA, Dunnett test). D, Summary of the rise time (10-90%) of the outward current in exWT and TPD receptors. The rise increased steadly in exWT receptors (p=0.005). The rise time was no significant change in the rise time in recordings from the TPD receptors (p>0.05).

## Discussion

The acute activation, desensitization, endocytosis and the development of tolerance to opioids were characterized in locus coeruleus neurons from a MOR knockout rat following the viral expression of exWT and TPD MORs. The results with the exWT receptors in the knockout animal were similar to those found in wild type animals. The activation kinetics of TPD receptors was also the same as both virally expressed and endogenous WT receptors. This study demonstrates that the elimination of phosphorylation of sites along the C-terminal of the MOR largely eliminates acute desensitization and the development of long-term cellular tolerance to morphine. The results also indicate that with the expression of receptors specifically in the locus coeruleus the development of tolerance is cell autonomous, independent of receptor activation in other areas of the CNS.

### Desensitization and internalization

MOR desensitization and internalization are not closely linked. The strongest evidence is based on experiments using mutation of the STANT sequence. Although mutations in the STANT sequence resulted in a significant decrease in internalization, acute desensitization was little changed (Lau et al., 2011, Just et al., 2013; Birdsong et al., 2015). The conclusion was that acute desensitization precedes internalization and the two processes are mechanistically distinct. The same conclusion was reached in experiments in cultured locus coeruleus neurons from a transgenic mouse that expressed soluble GFP under the control of the tyrosine hydroxylase promotor (TH-GFP). Internalization of the endogenous MORs measured with the use of a fluorescent peptide, dermorphin-Alexa594, was blocked by concanavalin A and yet desensitization and the recovery from desensitization measured electrophysiologically were not changed (Arttamangkul et al., 2006). The sequence beginning at T354 and ending at T357 (TSST) was also efficiently phosphorylated following the application of potent opioid agonists (Lau et al., 2011). An alanine-mutant of C-terminal phosphorylation sites excluding TSST and T394 (6S/T-A sites) prevented receptor endocytosis while desensitization remained unchanged (Yousuf et al., 2015). It was not until the TSST sequence along with the STANT sequence or all phosphosites on the C terminus were mutated to alanine that there was a large decrease in the degree of acute desensitization (Birdsong et al., 2015, Yousuf et al., 2015). Similar results were shown in this study with viral expressed TPD in MOR knockout rats. Thus it is clear that although acute desensitization and internalization of MOR are dependent on phosphorylation, the two processes involve mechanisms that can be distinguished by the degree of phosphorylation of the C-terminal tail.

The heterologous desensitization induced by muscarinic receptor activation was not blocked after mutations of C-terminal phosphorylation sites. Activation of muscarinic receptors can increase PKC activity that is thought to underlie heterologous desensitization of MORs (Bailey et al., 2004). The muscarine-induced decrease in activation of potassium conductance in slices expressing the TPD was similar to results obtained in slices from wild type animals (Shen & North, 1992; Fiorillo & Williams, 1996). The results may be interpreted in two ways. One is that the facilitation of MOR desensitization by PKC results from phosphorylation of intracellular loops of MOR and not at the carboxy tail (Chen et al., 2013). It is also possible that phosphorylation of other signaling proteins results in the inhibition (Chu et al., 2010). It has been shown that activation of muscarinic receptor alters the trafficking of MOR heterologous expressed in HEK293 cells (Lopez-Gimenez et al., 2017), however a previous trafficking study examining FLAG-MORs in the mouse locus coeruleus found that muscarine caused no change in internalization (Arttamangkul et al., 2015).

### Is the TPD a G protein biased receptor?

Depending on the agonist, MORs can activate differential downstream processes. A number of biased agonists have been described recently (Siuda et al., 2017). G-protein biased agonists have reduced ability to recruit arrestin while maintaining signaling through G proteins. Mutations in the C-terminus of MORs have a significant impact on arrestin signaling. Mutations of even one phosphorylation site at S375 and/or the STANT sequence in the C-terminus resulted in a significant reduction in the association with arrestin and the induction of internalization (Lau et al., 2011). The STANT mutant MOR was functional as measured by adenylyl cyclase (Leu et al., 2011) and activation of potassium conductance (Birdsong et al., 2015) assays. Mutations of all phosphorylation sites on the C-terminus (TPD) eliminated endocytosis induced by highly efficacious agonists such as DAMGO, etonitazene and fentanyl in HEK293 cells (Just et al., 2013) as well as ME in LC neurons (present study). Given these observations, the TPD receptor could be considered to function almost exclusively through a G protein-dependent process.

### Cellular tolerance and homeostatic mechanisms

One robust measure of tolerance in the LC is a decrease in the recovery from acute desensitization in slices from morphine treated animals (Dang & Williams, 2004; Quillinan et al, 2011; Levitt & Williams, 2012). In experiments from slices expressing TPD receptors, chronic morphine treatment had no effect. There are two possible conclusions that follow this observation. First, the near complete block of desensitization alone results in reduced tolerance. This however requires further investigation using a longer duration of morphine treatment. It is possible that tolerance may develop slower. It is also possible that compensatory homeostatic mechanisms due to continuous signaling may develop from the longer-term chronic treatment. The second conclusion centers on the G protein biased nature of this receptor and the elimination of arrestin binding leading to internalization. It may be that the activation of the arrestin pathway is a key step in the development of long-term tolerance. Studies using arrestin3-null mice show that cellular tolerance is attenuated (Quillinan et al., 2011; Dang et al., 2011, Connor et al., 2015). This could be the result of MOR trafficking because while the internalization of MORs in these animals was normal, receptor recycling was faster after chronic morphine treatment (Quillinan et al., 2011). However, interpretation of these studies is difficult given that only one isoform of arrestin was knocked out. Experiments expressing phosphorylation mutants in the C-terminus in the MOR knockout rat could begin to address the role of internalization in the development of cellular tolerance.

### Summary

The present work addresses one mechanism that underlies the development of long-term tolerance to morphine. Phosphorylation of the C-terminal of the MOR has been shown to be a key step in the mechanism of acute desensitization. Elimination of phosphorylation sites on the C-terminal rendered MORs resistant to cellular measures of long-term tolerance induced by morphine. One conclusion is that desensitization and/or internalization of MORs is necessary for the development of cellular tolerance to opioids. Although there were no obvious change in cellular excitability, it could be that continued signaling through the phosphorylation deficient receptors result in downstream homeostatic mechanisms that counteract the lack of cellular tolerance and may well increase signs of withdrawal.

## Materials and Methods

### Drugs

Morphine sulfate and morphine alkaloid were obtained from the National Institute on Drug Abuse (NIDA), Neuroscience Center (Bethesda, MD). Naloxone was purchased from Abcam (Cambridge, MA), MK-801, from Hello Bio (Princeton, NJ), UK14304 tartrate, from Tocris (Bio-Techne Corp. Minneapolis, MN). Potassium methanesulfonate was from Alfa Aesar (Ward Hill, MA). [Met^5^] enkephalin (ME), endomorphin-1, muscarine, scopolamine, idazoxan and other reagents were from Sigma (St. Louis, MO). Caged-enkephalin (CYLE) and Cagednaloxone (CNV-Nal) were a gift from Mathew Banghart.

Morphine alkaloid was converted to salt form with 0.1 M HCl and made up a stock solution in water. The working solution was diluted in artificial cerebrospinal fluid (ACSF) and applied during incubation or superfusion. Naloxone, endomorphin-1, muscarine, scopolamine, UK14304 tartrate and idazoxan were dissolved in water, diluted in ACSF and applied by bath superfusion. Bath perfusion of ME was with bestatin (10 mM) and thiorphan (1 mM) to limit breakdown of ME.

### Animals

All animal experiments were conducted in accordance with the National Institutes of Health guidelines and with approval from the Institutional Animal Care and Use Committee of the Oregon Health & Science University (Portland, OR). Adult (180 – 300 g or 5-6 weeks) male and female Sprague-Dawley rats were obtained from Charles River Laboratories (Wilmington, MA). A pair of MOR-knockout Sprague-Dawley rats with ZFN target site (GCTGTCTGCCACCCAgtcaaaGCCCTGGATTTC within exon 2) were generated by Horizon (St. Louis, MO) and received as F3 generation. The animals were bred and raised in house for two more generations before used in the experiments. The gene deletion was confirmed by genotyping using the primer 5’CATATTCACCCTCTGCACCA3’.

### Microinjection protocol

MOR-knockout animals (24-30 days) were anesthetized with isofluorane (Terrell^®^, Piramal Clinical Care, Inc., Bethlehem, PA) and placed in a stereotaxic frame for micro-injection of viral particles containing adeno associated virus type 2 for the expression of wild type MORs (exWT, AAV2-CAG-SS-GFP-MOR-WT-WPRE-SV40pA, 2.06x10^13^ vg/ml) and total phosphorylation deficient MORs (TPD, AAV2-CAG-SS-GFP-MOR-TPD-WPRESV40pA, 2.21x10^13^ vg/ml). Both viruses were obtained from Virovek (Hayward, CA). Injections of 200 nl at the rate of 0.1 μl/min were done bilaterally at ±1.25 mm lateral from the midline and -9.72 mm from the bregma at a depth of 6.95 mm from the top of the skull using computer controlled stereotaxic Neurostar (Kähnerweg, Germany). Experiments were carried out 2-4 weeks following the injection.

### Animal treatment protocols

Rats (5-6 weeks) were treated with morphine sulfate continuously released from osmotic pumps as described previously (Quillinan et al., 2011). Osmotic pumps (2ML1, Alzet, Cupertino, CA) were filled with the required concentration of morphine sulfate in water to deliver 80 mg/kg/day. Each pump has a 2 ml reservoir that releases 10 μl/hour for up to 7 days. Rats were anesthetized with isoflurane and an incision was made in the mid-scapular region for subcutaneous implantation of osmotic pumps. Pumps remained in animals until they were used for experiments 6 or 7 days later.

### Tissue preparation

Horizontal slices containing locus coeruleus (LC) neurons were prepared as described previously (Williams and North, 1984). Briefly, rats were killed and the brain was removed, blocked and mounted in a vibratome chamber (VT 1200S, Leica, Nussloch, Germany). Horizontal slices (250-300 μm) were prepared in warm (34°C) artificial cerebrospinal fluid (ACSF, in mM): 126 NaCl, 2.5 KCl, 1.2 MgCl_2_, 2.6 CaCl_2_, 1.2 NaH_2_PO_4_, 11 D-glucose and 21.4 NaHCO_3_126 and 0.01 (+) MK-801 (equilibrated with 95% O2/ 5% CO2, Matheson, Basking Ridge, NJ). Slices were kept in solution with (+)MK-801 for at least 30 min and then stored in glass vials with oxygenated (95% O2/ 5% CO2) ACSF at 34°C until used.

### Electrophysiology

Slices were hemisected and transferred to the recording chamber which was superfused with 34°C ACSF at a rate of 1.5 - 2 ml/min. Whole-cell recordings were made from LC neurons with an Axopatch-1D amplifier in voltage-clamp mode (V_hold_ = -60 mV). Recording pipettes (1.7 – 2.1 MΩ, World Precision Instruments, Saratosa, FL) were filled with internal solution containing (in mM): 115 potassium methanesulfonate or potassium methyl sulfate, 20 KCl, 1.5 MgCl_2_, 5 HEPES(K), 10 BAPTA, 2 Mg-ATP, 0.2 Na-GTP, pH 7.4, 275-280 mOsM. Series resistance was monitored without compensation and remained < 15 MΩ for inclusion. Data were collected at 400 Hz with PowerLab (Chart version 5.4.2; AD Instruments, Colorado Springs, CO). Intracellular recordings of membrane potential were made with glass electrodes (50-80 MΩ, World Precision Instruments, Saratosa, FL,) filled with KCl (2 M) and an Axoclamp-2A amplifier. Hyperpolarizing current (<20 pA) was used to prevent spontaneous firing of LC neurons. The depolarization induced by muscarine was corrected with the addition of more hyperpolarizing current to inhibit firing. Most drugs were applied by bath superfusion. In some experiments, [Leu^5^]enkephalin was applied by photolysis of caged-[Leu^5^]enkephalin (CYLE). A solution containing CYLE (30 μM), bestatin (1 μM) and thiorphan (10 μM) was recycled for photolysis experiments. In other experiments naloxone was released from the solution of caged-naloxone (CNV-Nal, 5 μM) recycled in the presence of agonist (ME 1 μM, or endomorphin-1 100 nM). Photolysis was carried out with a full-field illumination of a 365-nm LED lamp (Thorlabs, Inc., Newton, NJ) attached to the epifluorescence port.

### Anti-GFP nanobody expression and purification

A nanobody recognizing GFP was obtained from Addgene (Cambridge, MA) and cloned into the pET-22b vector with N-terminal 8xHis-tag followed by thrombin cleavage site. The lysine at 116 of nanobody was mutated to cysteine for a single dye-labeling site. Protein expression was conducted in *E-coli* strain BL21 (New England BioLabs, Ipswich, MA) in Terrific Broth medium. The cell culture was grown to OD_600_ 0.7 to 1.0 at 37°C, and protein synthesis was induced by 0.5 mM of isopropyl β-D-1-thiogalactopyranoside and was fermented at 20°C for 18 h. Cells were harvested by centrifugation and then lysed in a lysing buffer (in mM): 50 HEPES, 500 NaCl, 5 DL-dithiothretiol, 10 imidazol and 10% glycerol using a sonicator (10 min, 4 s sonication, 8 s paused, on ice). The debris was eliminated by centrifugation and the clear lysate was purified using the HisTrap column (GE Healthcare, Marlborough, PA). The His-tag was removed by adding thrombin protease (Sigma-Aldrich (St. Louis, MO) in to the protein solution at 1:100 (by mass) and incubated at 4°C overnight. The protein was further purified by size-exclusive chromatography (Superdex 200) in Dulbecco’s Phosphate Buffered Saline (Thermo Fisher Scientific, (Waltham, MA). Peak fractions having a single band by SDS-Page (10-20% gradient) electrophoresis were pooled and concentrated to ≈0.6mg/ml.

### Anti-GFP nanobody Alexa594 conjugation

The site-specific fluorescent labeling of the cysteine-mutated nanobody was modified from previously described (Pleiner et al., 2015). A solution containing the nanobody (100 μg) was used for the conjugation reaction. The solution was mixed with tris-(2-carboxyethyl)phosphine 15 mM on ice for 10 min. The buffer was exchanged to labeling buffer using P6 spin-column (BioRad, Hercules, CA). A labeling reaction was started by adding 1.5-fold of Alexa 594 maleimide (2 μl of 5 μg/μl in dimethylsulfoxide). The reaction proceeded on ice for 1 h. The conjugated nanobody was further purified by Superdex 200 in phosphate buffer. The degree of labeling was determined by measuring OD at 280 and 594 and was close to 1.

### MOR-GFP Trafficking

Brain slices (240 μm) from the virally injected rats were prepared as previously described. Slices were visualized with an Olympus Macroview fluorescent microscope for GFP expression in the LC area and then incubated in a solution of anti-GFP nanobody Alexa594 (Nb-A594, 10 μg/mL, 30-45 min). Images were captured with an upright microscope (Olympus, Center Valley, PA.) equipped with a custom-built two-photon apparatus and a 60x water immersion lens (Olympus LUMFI, NA1.1, Center Valley, PA). The dye was excited at 810 nm. Data were acquired and collected using Scan Image Software ((Pologruto et al., 2003). A z-series of 15-20 sections was collected at 1μm intervals. Drugs were applied by perfusion at the rate of 1 ml/min. All experiments were done at 35˚C. Internalization was calculated as percent of fluorescence in cytoplasm before and after ME application. The area of interest obtained by drawing the line along the plasma membrane.

### Data Analysis

Analysis was performed using GraphPad Prism 4 software (La Jolla, CA). Values are presented as mean ± SEM. Statistical comparisons were made using t-test or two-way ANOVA, as appropriate. Comparisons with p < 0.05 were considered significant.

## Author contributions

Conceptualization - SA & JTW, Tool building - SA, JRB, XS, Data Curation - SA, DAH, JTW, Formal Analysis, SA, DAH, JTW, writing-original draft – JTW, draft editing, SA.

## Animals/Key Resources

Al experiments were done in accordance with the Institutional Animal Care and Use Committees (IACUCs) at Oregon Health & Science University (OHSU). A colony of MOR knockout rats was maintained in house.

## Acknowledgements

This work was supported by NIH funding DA08163 (JTW). We thank members of the Gouaux lab for help in preparing the anti-GFP nanobody and members of the Williams lab for comments on the work.

## Figure legends

**Supplement Figure 1.**
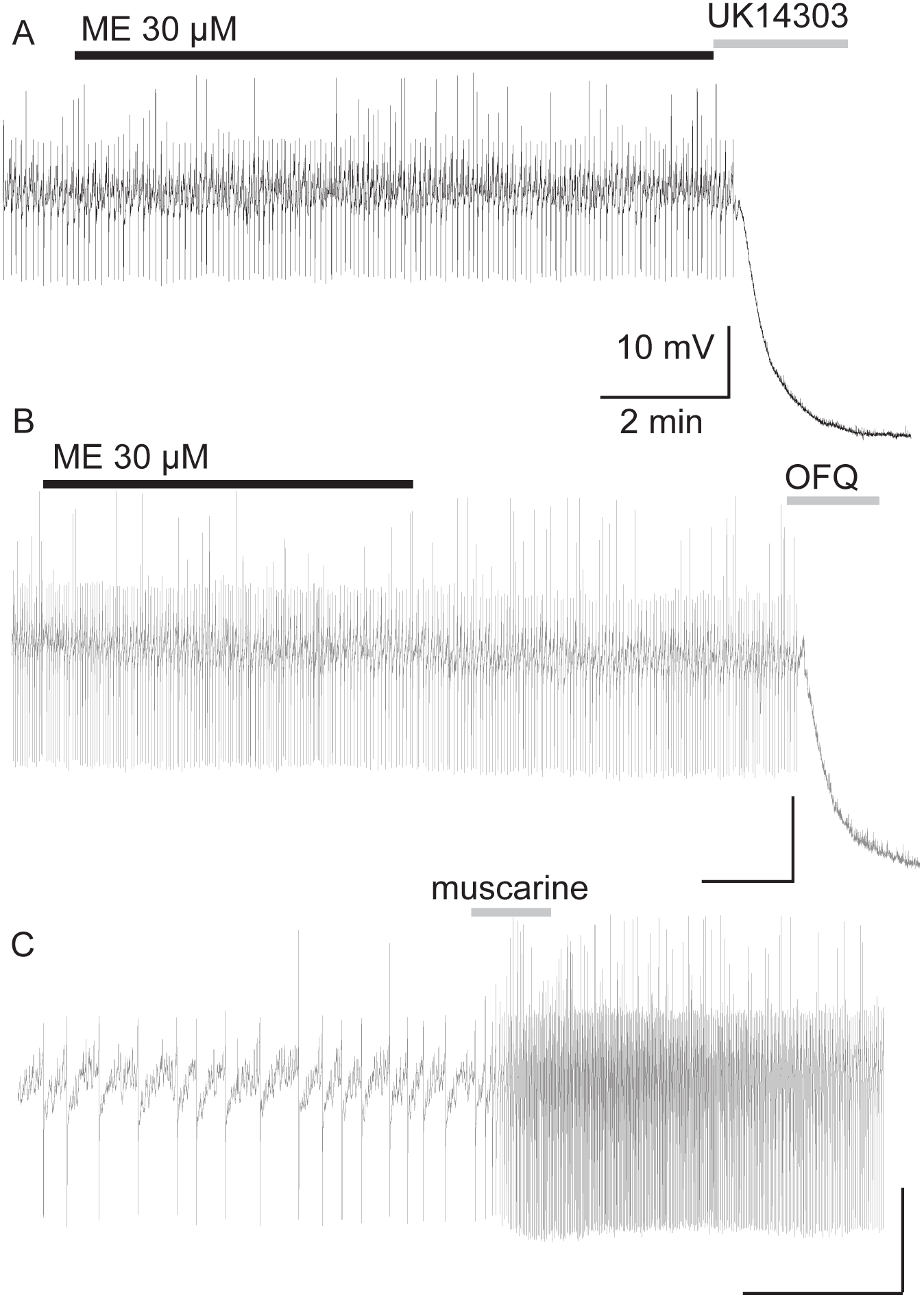
The firing rate of locus coeruleus neurons is not changed by opioids in recordings from the MOR knockout animal. Examples of recording with intracellular electrodes from three neurons. Application of ME (30 μM) had no effect on the firing rate however both UK14304 (3 μM,A) and OFQ (1 μM, B) inhibited firing and caused a hyperpolarization of the membrane potential. C, application of muscarine (10 μM) increased the firing rate of the LC neuron.

**Supplement Figure 4.**
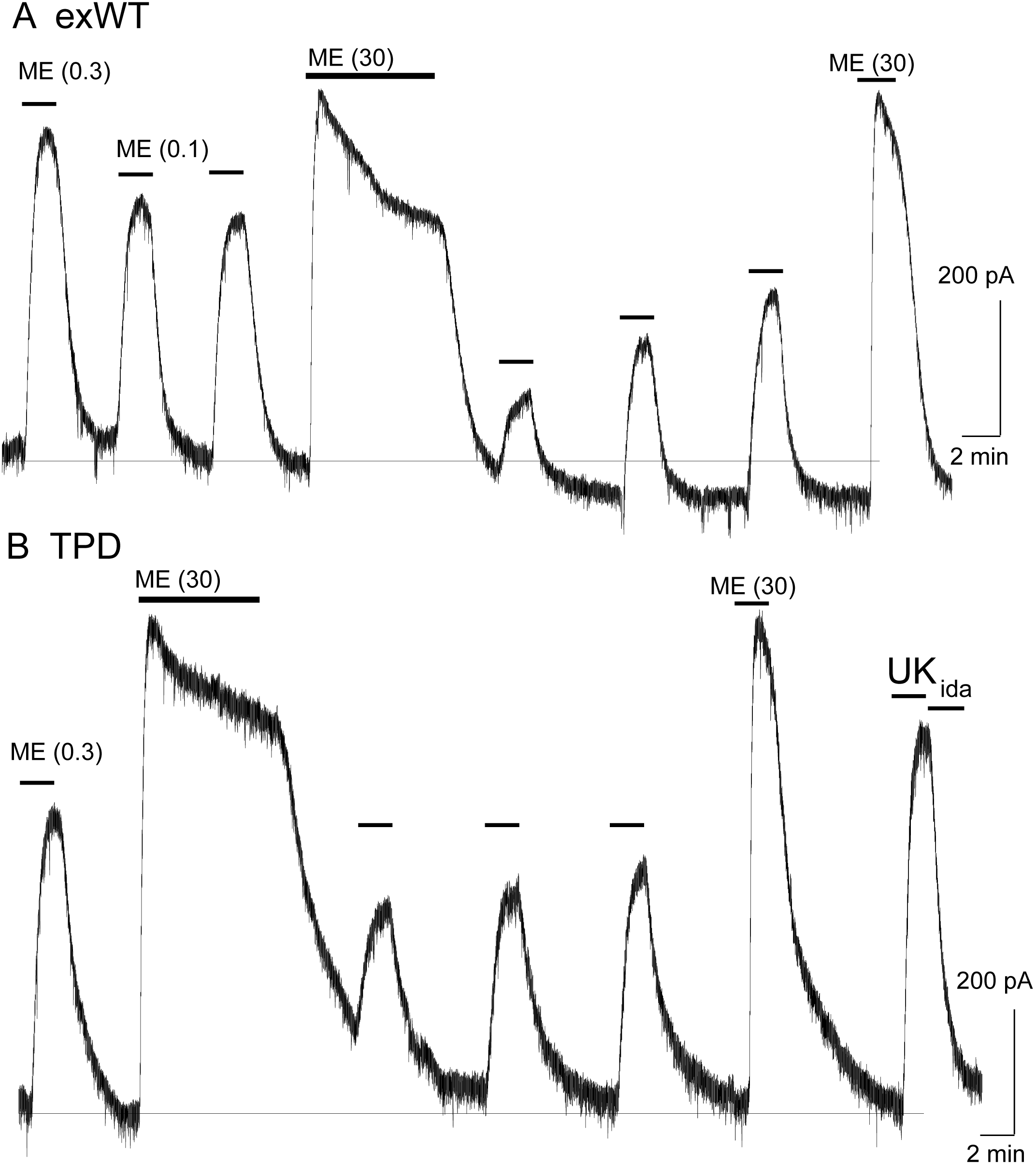
Reduced desensitization of the TPD receptor measured with whole cell voltage clamp recording. A, shows the outward current induced by application of different concentrations of ME. In this experiment ME (0.1 μM) was tested before and following the application of ME (30 μM, 10 min). The current induced by ME (30 μM) peaked and declined during the 10 min application and following the washout the current induced by ME (0.1 μM) was reduced and recovered slowly. B, the same experiment carried out with a neuron expressing the TPD receptor. The decline in the current induced by ME (30 μM) over 10 min was smaller than that in the exWT experiment and the decrease in the current induced by ME (0.3 μM) was also largely eliminated.

**Supplement Figure 5.**
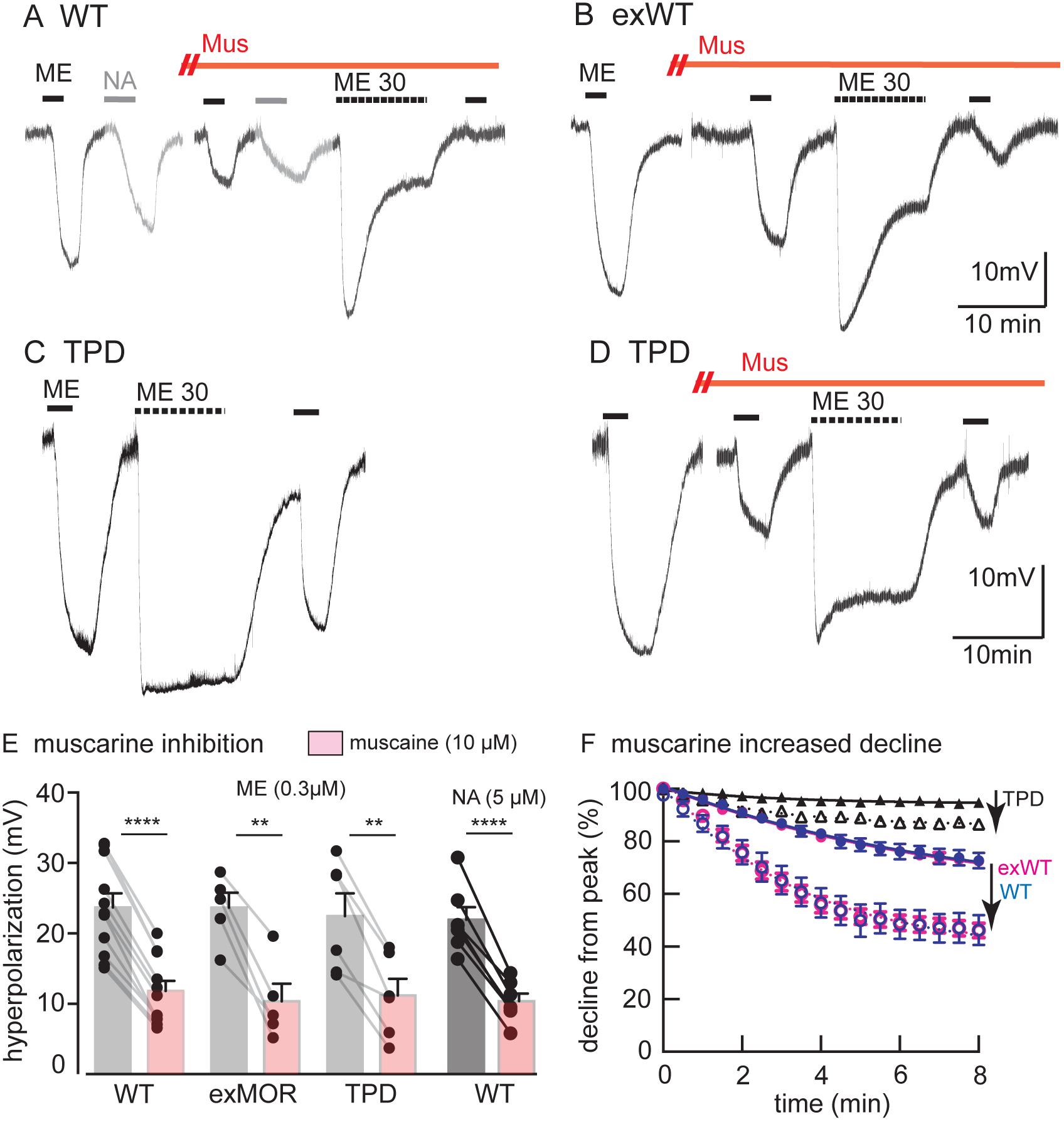
Whole cell voltage clamp experiment showing the inhibition of the outward current induced by photolysis of CYLE caused by muscarine (10 μM). A, an example of an experiment. Outward currents were induced before the application of muscarine (10 μM) and following the addition of the muscarinic antagonist, scopolamine (1 M). Muscarine caused in inward current and the outward current was reversibly decreased. B, the rate of rise of the current induced by photolysis of CYLE was reduced by muscarine, similar to that reported following acute desensitization. C, summarized results from experiments carried out in exWT and TPD receptors. Although desensitization was blocked in experiments with TPD receptors the inhibition by muscarine was not changed.

## References

Arttamangkul S, Birdsong W, Williams JT (2015) Does PKC activation increase the homologous desensitization of μ opioid receptors Br J Pharmacol 172: 583–592.

Arttamangkul S, Quillinan N, Low MJ, von Zastrow M, Pintar J, Williams JT (2008) Differential activation and trafficking of μ-opioid receptors in brain slices. Mol Pharmacol 74:972–979.

Arttamangkul S, Torrecilla M, Kobayashi K, Okano H, Williams JT (2006) Separation of mu-opioid receptor desensitization and internalization: endogenous receptors in primary neuronal cultures. J Neurosci 26: 4118–4125.

Bailey CP, Kelly E, Henderson G (2004). Protein kinase C activation enhance morphine-induced rapid desensitization of mu-opioid receptors in mature rat locus ceruleus neurons. Mol Pharmacol 66:1592–1598.

Birdsong WT, Arttamangkul S, Bunzow JR, Williams JT (2015) Agonist binding and desensitization of the μ-opioid receptor is modulated by phosphorylation of the C-terminal tail domain. Mol Pharmacol 88: 816–824.

Chen YJ, Oldfield S, Butcher AJ, Tobin AB, Saxena K Gurevich VV, Benovic JL, Henderson G, Kelly E (2013) Identification of phosphorylation sited in the COOH-terminal tail of the μ-opioid receptor J Neurochem 124, 189–199.

Chu J, Zheng H, Zhang Y, Loh HH, Law PY (2010) Agonist-dependent mu-opioid receptor signaling can lead to heterologous desensitization. Cell Signal 22: 684–696.

Connor M, Bagley EE, Chieng BC, Christie MJ (2015) β-Arrestin-2 knockout prevents development of cellular μ-opioid receptor tolerance but does not affect opioid-withdrawal-related adaptations in single PAG neurons. Br J Pharmacol 172: 492–500.

Dang VC, Chieng B, Azriel Y, Christie M (2011) Cellular morphine tolerance produced by β arrestin-2-dependent impairment of μ-opioid receptor resensitization. J Neurosci 31: 7122–7130.

Dang VC, Williams JT (2004) Chronic morphine treatment reduces recovery from opioid desensitization. J Neurosci 24: 7699–7706.

Feng B, Li Z, Wang JB (2011) Protein kinase C-mediated phosphorylation of the μ-opioid receptor and its effects on receptor signaling. Mol Pharmacol 79: 768–775.

Fiorillo CD, Williams JT (1996) Opioid desensitization: interactions with G-protein coupled receptors in the locus coeruleus. J Neurosci 16:1479–1485.

Illing S, Mann A, Schulz S (2014) Heterologous regulation of agonistindependent μ-opioid receptor phosphorylation by protein kinase C. Br J Pharmacol 171: 1330–1340.

Just S, Illing S, Trester-Zedlitz M, Lau EK, Kotowski SJ, Miess E, Mann A, Doll C, Trinidad JC, Burlingame AL, von Zastrow M, Schulz S (2013) Differentiation of opioid drug effects by hierachical multi-site phosphorylation. Mol Pharmcol 83: 633–639.

Lau EK, Trester-Zedlitz M, Trinidad JC, Kotowski SJ, Krutchinsky AN, Burlingame AL, von Zastrow M (2011) Quantitative encoding of the effect of a partial agonist on individual opioid receptors by multisite phosphorylation and threshold detection. Sci Signal 4: ra52.

Levitt ES, Williams JT (2012) Morphine desensitization and cellular tolerance are distinguished in rat locus ceruleus neurons. Mol Pharmacol 82: 983–992.

Lopez-Gimenez JF, Alvarez-Curto E, Milligan G (2017) M3 muscarinic acetylcholine receptor facilitates the endocytosis of mu opioid receptor mediated by morphine independently of the formation of heteromeric complexes. Cell Signal 35: 208–222.

Pleiner T, Bates M, Trakhanov S, Lee CT, Schliep JE, Chug H, Böhning M, Stark H, Urlaub H, Görlich D (2015) Nanobodies: site-specific labeling for super-resolution imaging, rapid epitope-mapping and native protein complex isolation. Elife 4: e11349.

Pologruto TA, Sabatini BL, Svoboda K (2003). Scanimage: flexible software for operating laser scanning microscopes. Biomed Eng Online 2:13.

Quillinan N, Lau EK, Virk M, von Zastrow M, Williams JT (2011) Recovery from μ-opioid receptor desensitization after chronic treatment with morphine and methadone. J Neurosci 31: 4434–4443.

Shen KZ, North RA (1992) Muscarine increases cation conductance and decreases potassium conductance in rat locus coeruleus neurons. J Physiol 455: 471–485.

Wang HL, Chang WT, Hau PC, Chow YW, Li AH (2002) Identification of two C-terminal amino acids, Ser(355) and Thr(357), required for short-term homologous desensitization of mu-opioid receptors. Biochem Pharmacol 64: 257–266.

Williams JT, Ingram SL, Henderson G, Chavkin C, von Zastrow M, Schulz S, Koch T, Evans CJ, Christie MJ (2013) Regulation of μ-opioid receptors: desensitization, phosphorylation, internalization, and tolerance. Pharmacol Rev 65:223–254.

Williams JT, North RA (1984) Opiate-receptor interactions on single locus coeruleus neurons. Mol Pharmacol 26: 489–497.

Yousuf A, Miess E, Sianati S, Du YP, Schulz S, Christie MJ (2015) Role of phosphorylation sites in desensitization of μ-opioid receptor. Mol Pharmacol 88: 825–835.

